# Biochemical and Phytochemical Efficacy of Defatted *Moringa oleifera* Seeds in Alleviating Protein-Energy Malnutrition

**DOI:** 10.1101/2024.10.14.618128

**Authors:** Raphael Eneji Jegede, Gideon Ayeni, Rose Mafo Abaniwo, Godwin Amoka Audu, Abdullahi Haruna

## Abstract

Protein-energy malnutrition (PEM) is a severe health condition affecting millions, especially in developing regions. This study investigates the potential of *Moringa oleifera* seeds as a low-cost protein source to address protein energy malnutrition. *Moringa oleifera* seeds were defatted using different solvents (n-hexane, acetone, and a mixture of n-hexane and acetone) and used in formulating diets for PEM-induced Wistar rats. The study analyzed the seeds’ phytochemical constituents, proximate composition, amino acid profiles, and bioactive compounds through Gas Chromatograph-Mass Spectroscopy (GC-MS). Twenty (2) wistar Rats were randomly assigned to four groups: Group A, control group, B= PEM-induced group, Group C, malnourished rats treated with 15% soya bean based-diet and group D, malnourished rats treated with 15% defatted *Moringa oleifera* seed-based diet. The results show that defatting increased protein content and reduced anti-nutritional factors like oxalates, saponins, and tannins, enhancing nutrient bioavailability. Defatted *Moringa oleifera* seed-based diets improved liver and kidney functions, lipid profiles, and protein digestibility in protein energy malnutrition-induced rats. Additionally, the seeds contained essential bioactive compounds with potential antioxidant, antimicrobial, and anti-inflammatory properties. These findings suggest that *Moringa oleifera* seeds could be a promising alternative protein source for combating protein energy malnutrition.

---

Protein-energy malnutrition (PEM) is a severe health condition that arises due to inadequate intake of protein and energy, affecting millions, particularly in low-income and developing countries(Naheed *et al*., 2021). PEM is particularly common among children and can result in conditions such as kwashiorkor, marasmus, or marasmic-kwashiorkor, a combination of both. It leads to impaired growth, weakened immune function, and a higher susceptibility to infections, contributing significantly to child mortality (Michael *et al*., 2022). According to the World Health Organization (WHO), malnutrition effects nearly half of all child deaths globally (WHO, 2020).

PEM can develop due to various factors, including poor dietary intake, infections, and socioeconomic conditions. While the primary cause is inadequate nutrient intake, the condition can also be aggravated by diseases like diarrhea, which reduces the body’s capacity to absorb nutrients (Mathewson *et al*., 2021). Children, pregnant women, and the elderly are most vulnerable to PEM due to their increased nutritional needs (Rasheed *et al*., 2023).

The management of PEM involves dietary interventions aimed at restoring nutritional balance by providing energy and protein in adequate amounts. Typically, treatment includes the use of ready-to-use therapeutic foods (RUTFs) which are energy-dense, micronutrient-enriched foods used in treating severe malnutrition(Okhiria, 2007). However, the high cost of RUTFs can limit their accessibility, particularly in resource-poor regions, highlighting the need for affordable and locally available alternatives.

Traditional therapies for PEM include dietary formulations based on animal proteins, such as milk and eggs, as well as plant-based protein sources. However, due to high cost of animal protein and soya bean, these options may not be accessible in some regions, prompting the search for alternative protein-rich foods, such as *Moringa oleifera*(Awuchi *et al*., 2020).

*Moringa oleifera*, commonly known as the drumstick tree, is widely recognized for its exceptional nutritional and medicinal properties. Native to parts of South Asia and Africa, the tree has gained international recognition due to its ability to thrive in harsh climatic conditions and provide essential nutrients in regions with food insecurity (Gopalakrishnan *et al*., 2016)). *Moringa oleifera* seeds are especially rich in protein, vitamins, minerals, and bioactive compounds, making them a viable option for addressing malnutrition(Dhakar *et al*., 2011).

The seeds of *Moringa oleifera* are an excellent source of protein, containing all the essential amino acids required by the body. According to research, Moringa seeds contain approximately 30-35% protein, comparable to commonly consumed legumes like soybeans (Gopalakrishnan *et al*., 2016). In addition to its protein content, the seeds are rich in vitamins (A, C, E), minerals (calcium, magnesium, potassium), and antioxidants. These nutritional attributes make *Moringa oleifera* seeds an ideal supplement in diets for treating PEM(Ayodele *et al*., 2021). Moreover, Moringa seeds possess a balanced amino acid profile, which is essential for the growth and repair of tissues, making it particularly suitable for combating protein deficiency in PEM (Dhakar *et al*., 2011). Additionally, Moringa seeds are a good source of fats and essential fatty acids, which are crucial in providing energy and promoting cognitive development in malnourished children (Ashraf & Gilani, 2007).

## Materials and methods

### Chemicals and Reagents

All reagents used are of analytical grade and were procured from BDH, UK.

### 2.2. Plant Materials and Authentication

*Moringa oleifera* seeds were obtained from the premises of Faculty of Agriculture, Prince Abubakar Audu University, Anyigba, Kogi state. The seeds were identified and authenticated at the herbarium unit, Plant Biology Department, University of Ilorin, Ilorin, Nigeria with specimen voucher number (UILH/001/1585/2023) deposited.

### Defatting of *Moringa oleifera* kernel

*The seed coat of M. oleifera* seeds were removed, the kernel oven dried (SM 9053, Surgifriend Medicals, England) at 40°C to a constant weight, which was then pulverized with electric blender (Master Chef, Model MC-BL 1644, Jiangsu, China). 200g of the pulverized *M. oleifera* kernels was weighed into a dry conical flask. 800mls of n-hexane was added and the flask shaken vigorously. It was then left for 24hrs for the lipid to extract from the kernel. The solvent containing the extracted lipid was then decanted off and filtered using Whatman filter paper no 1. The process was repeated for 2 more times to improve the efficiency of the defatting process (Jegede *et al*., 2020). After the solvent has been filtered, the residue was spread on aluminum foil at room temperature to evaporate the residual solvent and dried the residue to fine powder and packed into an air tight container for feed formulation and analyses.

### 2.3. Experimental Animals

Twenty (20) White Albino rats of wistar strain [average weight ±SEM (53.21±2.74g) were procured from animal holding unit of the department of biochemistry, Prince Abubakar Audu University, Anyigba, Kogi State, Nigeria for the research. The rats were housed in clean plastic cages located at a well-ventilated room, maintaining specific housing conditions: a 12-hour light-dark cycle, temperature ranging between 25–27°C, and humidity levels of 45%–55% (according to Weather.com). They were fed with pelletized diet from Chikun Feeds Limited, Ilorin, Nigeria, and they had continuous access to clean water and were acclimatized for a week before the commencement of the experiment.

### Induction of Protein Energy Malnutrition

Protein Energy Malnutrition (PEM) was induced in fifteen (15) rats by feeding them with low protein isocaloric diet *ad libitum* and clean water for 28days according to modified method of Adelusi & Olowokere, (1985). The composition of the diet is as shown in Table 1.

**Table 1:** Composition of Low protein Isocaloric Diet.

| Composition | Low protein Isocaloric diet (g/Kg) |
| --- | --- |
| Soya Bean | 40g |
| Corn Starch | 516g |
| Sucrose | 300g |
| Cellulose | 40g |
| Soya Bean oil | 50g |
| DL-methionine | 4g |
| * Vitaflash Amino WSP | 50g |
| Total | 1000g |

\*VITAFLASH AMINO WSP (vitamin-Amino Acids):vitamin A 10,000,000 IU; vitamin D3 2,000,000 IU; vitamin E 15,000 mg; vitamin K3 2,500mg; vitamin B1 1,000mg; vitamin B2 2,000mg; vitamin B6 2,000mg; vitamin B12 10,000mg; folic acid 300mg; Ca-d-pantothenate 7,500mg; nicotine acid 20,000mg; choline chloride 15,000mg; vitamin C 40,000mg;DL-Methionine 50,000mg; L-Lysine 50,000mg; amino acids 52,000mg (cysteine,tryptophan,arginine,threonine,isoleucine,leucine,valine,histidine,phenylalanine,tyrosine and glycine).

### Animal Grouping

Twenty (20) wistar rats were randomly assigned into four groups each containing five (5) rats: A, B, C and D. group A was fed with rat feed (Chikun Feeds, Ilorin, Nigeria) and water *ad libitum* and group B, C and D, were fed with low protein isocaloric diet (Table 1) and clean water *ad libitum* for 28days to induce protein energy malnutrition. On the 29^th^ day, animals in group A and B were sacrificed while animals in group C and D were fed with various treatment diets (Table 2) for another 28days and on the 29^th^ day they were sacrificed.

**Table 2:** Composition of Treatment diets.

| Composition | 15%Soya Bean Protein (g/Kg) | 15% *M. oleifera* kernel Protein based-diet (g/Kg) |
| --- | --- | --- |
| Soya Bean | 150 | - |
| *M. oleifera* | - | 150 |
| Corn Starch | 406 | 406 |
| Sucrose | 300 | 300 |
| Cellulose | 40 | 40 |
| Soya Bean oil | 50 | 50 |
| DL-methionine | 4 | 4 |
| * Vitaflash Amino WSP | 50 | 50 |
| Total | 1000 | 1000 |

VITAFLASH AMINO WSP (vitamin-Amino Acids):vitamin A 10,000,000 IU; vitamin D3 2,000,000 IU; vitamin E 15,000 mg; vitamin K3 2,500mg; vitamin B1 1,000mg; vitamin B2 2,000mg; vitamin B6 2,000mg; vitamin B12 10,000mg; folic acid 300mg; Ca-d-pantothenate 7,500mg; nicotine acid 20,000mg; choline chloride 15,000mg; vitamin C 40,000mg; DL-Methionine 50,000mg; L-Lysine 50,000mg; amino acids 52,000mg (cysteine,tryptophan,arginine,threonine,isoleucine,leucine,valine,histidine,phenylalanine,tyrosine and glycine).

A: Control Rats fed with commercial diet (Naïve)

B: Rats fed with Low Protein isocaloric diet only

C: Rats fed with Low Protein Isocaloric diet but treated with 15% soya bean Based Diet

D: Rats fed with Low Protein isocaloric diet but treated with 15% defatted *M. oleifera* kernel based diet

### Animal sacrifice

On the 29th day post treatment, animals were sacrificed according to the method described by Yakubu *et al*., (2005) after they were anaesthetized by diethyl ether and blood was collected by cardiac puncture.

### Determination of body weight

Body weight measurement was done using 0.01g sensitive weighing balance.

### Quantitative Phytochemical Constituents and Nutritional Composition of *M. oleifera* kernel Determination

The alkaloid content was quantified following the procedure outlined by Adeniyi *et al*., (2012), The saponin (Obadoni & Ochuko, 2002), phenolic content ( Djurdjević *et al*., 2007), flavonoid (Boham & Kocipai-Abyazan, 1974; Harborne & Mabry, 2013), tannin and cyanogenic glycosides (Ibukun *et al*., 2013), oxalate content was estimated according to the modified method of (Karamad *et al*., 2019), Phytic acid (Harland *et al*., 1986), Mineral elements were determined following the standard procedures of the Association of Official Analytical Chemists (AOAC) (Poitevin, 2016), amino acids profile was determine following the method described by AOAC with modifications (Otemuyiwa & Adewusi, 2013), The GC-MS analysis (Dowd *et al*., 2010), The proximate compositions were determined in accordance with AOAC (Horwitz & Latimer, 2005), The carbohydrate was determined by calculation using the formula:

Carbohydrate = 100 - (Crude protein +Fat +Ash + Crude fibre + Moisture)

*In-vitro* Protein digestibility was carried out following the method developed by Mertz *et al*., (1984) with modifications as reported by Gulati *et al*., (2017).

### Biochemical and Enzymes Assays Determination of Plasma Liver Function Indices

The plasma enzymes activity were determined spectrophotometrically using commercial assay kits, Aspartate amino transferase and Alanine amino transferase were quantified following procedure outlined by Reitman & Frankel, (1957), Alkaline Phosphatase (Ahlers, 1975), total protein was determined by biuret method as (Gornall *et al*., 1949), Albumin was quantified as described by (Doumas, 1975), globulin (Goldenberg & Drewes, 1971), total and conjugated bilirubin (Otsuji *et al*., 1988).

### Determination of Kidney Function Indices

The plasma electrolytes were estimated using flame photometer and Atomic Absorption Spectrophotometer (AAS). Serum sodium was quantified according to the method of Levy, (1981), Potassium and chloride ion (Steffes & Freier, 1976), calcium (Busch *et al*., 1974), and bicarbonate (Jørgensen & Astrup, 1957). The plasma urea concentration was determined by the procedure of Veniamin & Vakirtzi-Lemonias, (1970),and creatinine by Bartels *et al*., (1972).

### Determination of Plasma Lipid Profile

The plasma lipid profile was determine using commercial kits. High Density lipoprotein-cholesterol was determine spectrophotometrically by the method of Izzo *et al*., (1981), Low Density Lipoprotein LDL (Friedewald *et al*., 1972), Triglyceride (Mendez *et al*., 1975), Total cholesterol was estimated by the method of Allain *et al*., (1974) and the atherogenic indices (Kavitha & Sujatha, 2017).

## Results and Discussion

## Discussion

### Phytochemical constituents

The results of the phytochemical analyses is shown in Table 3. It revealed that defatting the kernel significantly (p < 0.05) all the phytochemicals constituents.

**Table 3:**
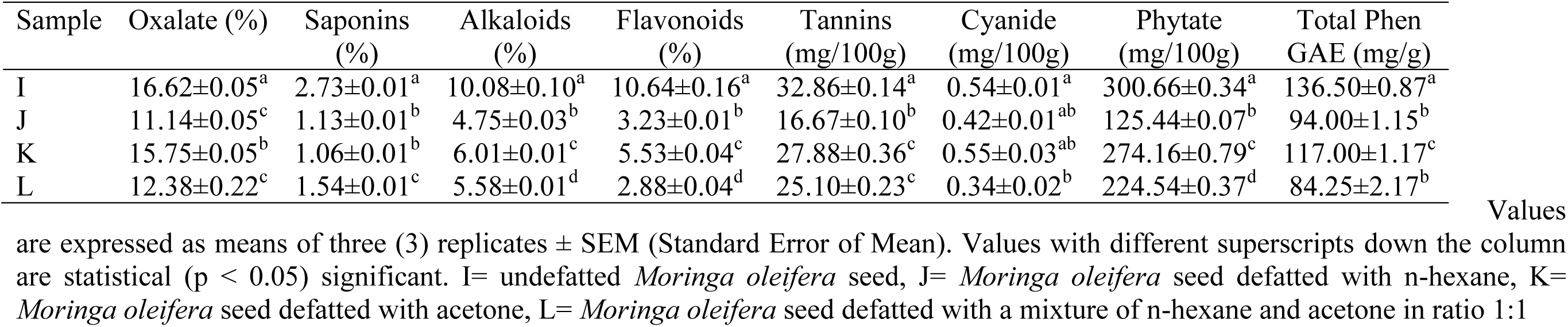
Phytochemical constituents of *M. oleifera* kernel and the defatted kernels.

Oxalates can bind to minerals like calcium, reducing their bioavailability. The study shows that defatting Moringa seeds with n-hexane (Sample J) resulted in the lowest oxalate content, while the highest was found in the undefatted seeds (Sample I). For protein energy malnourished patients, excessive oxalates could have a detrimental effect, as it may limit the absorption of essential minerals needed for recovery from malnutrition(Mihrete, 2019). Thus, defatted seeds, especially those treated with n-hexane, may be more beneficial for ensuring mineral availability. Saponins have potential to interfere with nutrient absorption and cause gastrointestinal issues in large amounts. Undefatted seeds had the highest saponin content, while the acetone-defatted (Sample K) and n-hexane-defatted seeds (Sample J) showed significantly (p < 0.05) lowered levels. Lowered saponin levels in defatted kernel make it more suitable for individuals suffering from PEM, as they may reduce the risk of nutrient malabsorption and gastrointestinal distress (Y. Zhang *et al*., 2023).

Alkaloids have medicinal properties but can be toxic in high amounts. Undefatted seeds had the highest alkaloid content, while the n-hexane-defatted seeds had the lowest. Alkaloids help to boast immune function, which is crucial for malnourished individuals (Olarewaju *et al*., 2022). However, a lower alkaloid concentration, as seen in defatted samples, may reduce potential toxic effects (Adibah & Azzreena, 2019).

Flavonoids, possess antioxidant properties and are vital in protecting against oxidative stress, which is often heightened in malnourished individuals. The defatting process though significantly (p < 0.05) reduce its concentration, but the quantity seen in n-hexane defatted form is sufficient in mopping out free radicals that might be generated during malnourishment. For PEM patients, the antioxidant capacity of the seeds might help mitigate inflammation and cell damage (Ahmed *et al*., 2022), so a balance needs to be considered in relation to their other benefits.

Tannins, though potentially beneficial as antioxidants, may interfere with the digestibility of proteins and iron absorption. Undefatted kernel contained the highest tannin levels, while defatting significantly (p < 0.05) reduce its levels. Lower tannin levels, as seen in the defatted samples, especially the n-hexane defatted sample might enhance protein digestibility and iron absorption in malnourished individuals, which is essential for recovery (J. Zhang *et al*., 2019).

The kernel contains a lowered levels of Cyanide content, a potentially harmful compound. The defatting process further significantly (p < 0.05) reduced its concentration. Since cyanide in high amounts can be toxic, the relatively low levels across all samples make them safer for consumption(Tahir *et al*., 2024).

Phytates can inhibit mineral absorption, which is critical for malnourished individuals who require increased mineral intake. The lowest phytate content was found in the n-hexane-defatted sample, while the highest was in the undefatted seeds. Defatting significantly (p < 0.05) reduces phytate levels, thereby increasing the bioavailability of essential minerals like iron, calcium, and zinc, which are crucial for malnutrition recovery(Grases *et al*., 2017).

Phenols have antioxidant properties, playing a protective role in the body. Undefatted seeds had the highest total phenol content, while defatting significantly (p < 0.05) reduced concentration, especially in Sample J. For PEM patients, phenols could help mitigate oxidative damage, though lower levels in defatted samples suggest that other nutritional benefits, such as improved protein and mineral absorption are considered alongside its antioxidant potential (Shahidi *et al*., 2019).

In addressing PEM, defatted *Moringa oleifera* seeds, especially the sample defatted with n-hexane, may offer better options due to lower levels of anti-nutritional factors such as oxalates, saponins, tannins, and phytates. These reductions enhance nutrient bioavailability, making these kernel more suitable for malnourished individuals (Kavi Kishor *et al*., 2021).

### Proximate compositions

The result of proximate composition is presented in table 4. The crude protein content significantly (p < 0.05) increased across the samples after defatting, with the highest value observed in Sample L, which was defatted with a mixture of n-hexane and acetone (1:1). This suggests that the defatting process enhances protein concentration by removing the fat content. Sample I (undefatted *Moringa oleifera* seed) showed a lower protein value, as the presence of fat dilutes the protein content. Defatting with n-hexane (Sample J) and acetone (Sample K) also showed increased protein levels, with significant differences compared to the undefatted sample (*p* < 0.05). This suggests that the type of solvent used in defatting affects the protein yield, with acetone defatting showing slightly lower protein content than n-hexane. Increased protein content is vital in addressing protein-energy malnutrition (PEM), as dietary protein is essential for growth, tissue repair, and overall metabolism (Wu, 2016).

**Table 4:** Proximate Compositions of *M. oleifera* kernel and the defatted kernels.

| Sample | Crude Protein (%) | Fat (%) | Ash (%) | Crude Fiber (%) | Moisture (%) | NFE (%) | Energy Value (kcal/100g) |
| --- | --- | --- | --- | --- | --- | --- | --- |
| I | 35.66±0.20<sup>a</sup> | 14.29±0.05<sup>a</sup> | 3.84±0.04<sup>a</sup> | 10.06±0.06<sup>a</sup> | 3.81±0.03<sup>a</sup> | 32.36±0.02<sup>a</sup> | 346.98±0.11<sup>a</sup> |
| J | 46.33±0.13<sup>b</sup> | 3.13±0.04<sup>b</sup> | 4.42±0.04<sup>b</sup> | 5.45±0.08<sup>b</sup> | 6.96±0.03<sup>b</sup> | 33.72±0.07<sup>b</sup> | 299.93±0.10<sup>b</sup> |
| K | 45.92±0.02<sup>b</sup> | 2.33±0.03<sup>c</sup> | 3.84±0.02<sup>a</sup> | 3.39±0.01<sup>c</sup> | 5.41±0.01<sup>c</sup> | 39.13±0.06<sup>c</sup> | 309.81±0.08<sup>c</sup> |
| L | 50.58±0.17<sup>c</sup> | 2.63±0.02<sup>d</sup> | 4.96±0.04<sup>c</sup> | 3.16±0.03<sup>d</sup> | 5.77±0.05<sup>d</sup> | 32.92±0.18<sup>ab</sup> | 312.43±0.14<sup>c</sup> |

Values are expressed as means of three (3) replicates ± SEM (Standard Error of Mean). Values with different superscripts down the column are statistical (p < 0.05) significant. I= undefatted *Moringa oleifera* seed, J= *Moringa oleifera* seed defatted with n-hexane, K= *Moringa oleifera* seed defatted with acetone, L= *Moringa oleifera* seed defatted with a mixture of n-hexane and acetone in ratio 1:1

The fat content was significantly (p < 0.05) reduced in all defatted samples compared to the undefatted Sample I, with Sample K (acetone-defatted) showing the lowest fat value (*p* < 0.05). Sample J (n-hexane) and Sample L (n-hexane + acetone) also had low fat content but were slightly higher than Sample K. The significant reduction in fat after defatting enhances the utility of Moringa seeds for protein supplementation in PEM treatment without contributing excess calories from fat, which can lead to metabolic disturbances (Lambe, 2019).

Ash content, which represents the mineral composition, was highest in Sample L (4.96±0.04%), indicating that the mixed solvent of n-hexane and acetone might extract fat more effectively, preserving and even enhancing the mineral content. Sample I (undefatted) and Sample K (acetone-defatted) showed similar ash content, while Sample J (n-hexane) showed a slight increase. Minerals play a significant role in addressing malnutrition, particularly in improving bone health, electrolyte balance, and enzymatic processes (Shankar, 2020).

The crude fiber content was significantly (p < 0.05) higher in Sample I compared to the defatted samples. This is expected, as defatting processes concentrate other macronutrients while reducing fiber content. Among the defatted samples, Sample J (n-hexane-defatted) had a higher fiber content than Sample K (acetone-defatted) and Sample L (mixed solvent), showing that solvent choice also influences fiber retention. Dietary fiber is essential in the management of malnutrition-related gastrointestinal issues, aiding digestion and improving nutrient absorption (Shozib *et al*., 2018).

Sample J (n-hexane) had the highest moisture content, followed by Sample L and Sample K. Sample I (undefatted) had the lowest moisture content, suggesting that defatting increases the moisture retention capacity of Moringa seeds. Higher moisture content may affect the shelf life and storage of the defatted seeds, but it can also improve the texture and palatability of food products formulated from these seeds (Anselm *et al*., 2023).

The nitrogen-free extract, representing the carbohydrate content, was highest in Sample, followed by Sample L and Sample J. Sample I (undefatted) had a similar NFE value to the defatted samples. Carbohydrates, while not as critical in addressing PEM as proteins, still provide essential energy for individuals suffering from malnutrition (Siva Kiran *et al*., 2015).

The energy values for each sample show that Sample I (undefatted) had the highest caloric content due to its higher fat content, which is more energy-dense than protein or carbohydrates. Sample J (n-hexane) had the lowest energy value, while Samples K and L had moderate energy values. The reduced caloric content in the defatted samples makes them ideal for protein supplementation in malnutrition without providing excess calories that could potentially lead to obesity or other metabolic conditions, especially in the management of protein-energy malnutrition (Siva Kiran *et al*., 2015).

### Mineral Element Compositions

The mineral element analysis of *Moringa oleifera* seeds is shown in figure 1. The result revealed Sodium (Na) as the most abundant element. Sodium is an essential mineral that plays a vital role in maintaining fluid balance, nerve signaling, and muscle function(Ali, 2023).

**Figure 1:**
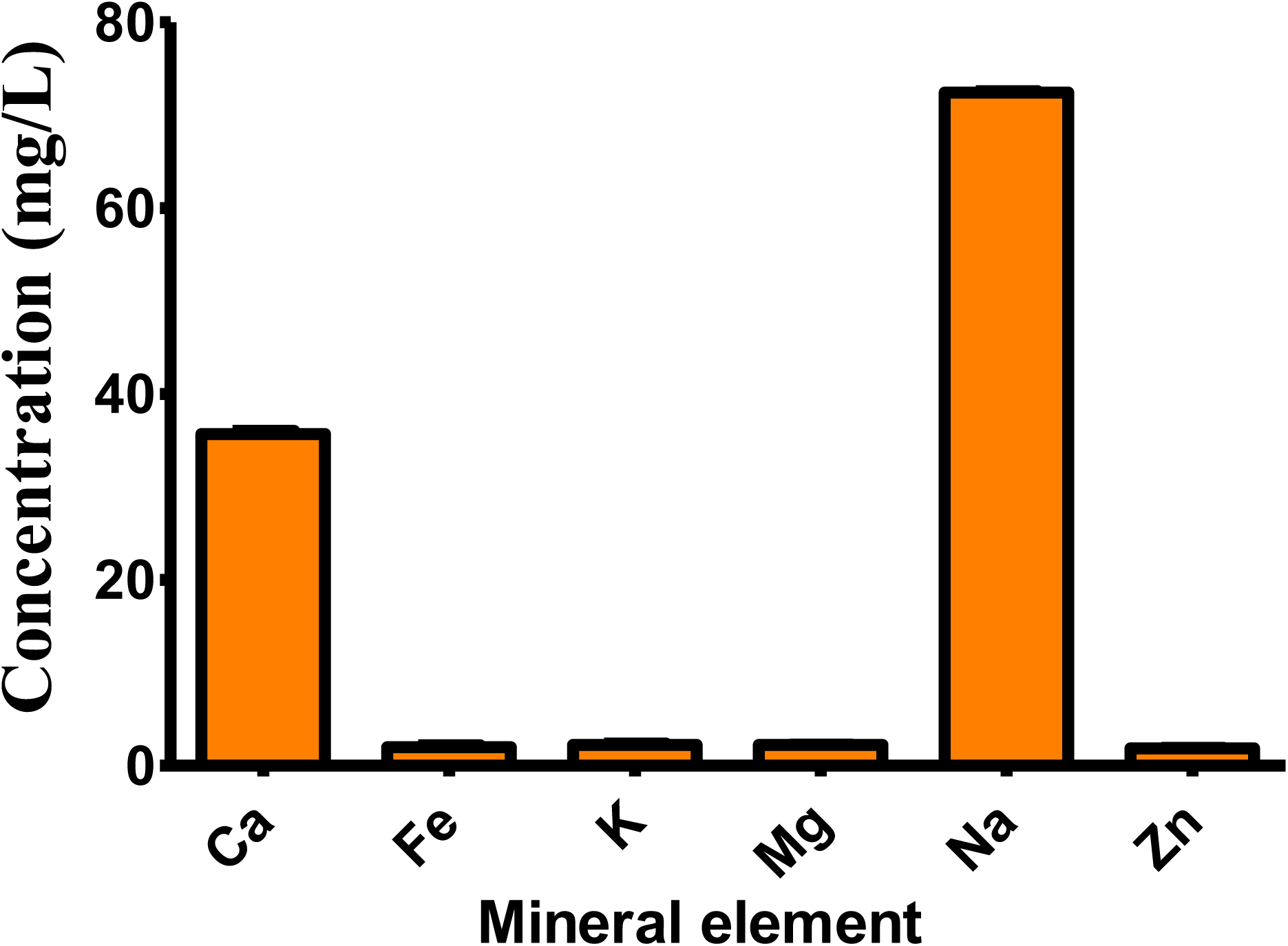
Mineral Elements of *M. oleifera* kernel and the defatted kernels. Values are expressed as means of three (3) replicates ± SEM (Standard Error of Mean). I= undefatted *Moringa oleifera* seed, J= *Moringa oleifera* seed defatted with n-hexane, K= *Moringa oleifera* seed defatted with acetone, L= *Moringa oleifera* seed defatted with a mixture of n-hexane and acetone in ratio 1:1.

Calcium (Ca) is the second most abundant mineral. Calcium is crucial for bone health, muscle contractions, and enzymatic processes. The relatively high concentration of calcium in *Moringa* seeds makes it an important contributor to nutritional intake, particularly in populations at risk of calcium deficiency(Kamran *et al*., 2020; Saa *et al*., 2019). On the lower end, zinc (Zn) was the least abundant element. Despite its low concentration, zinc remains essential for immune function, protein synthesis, and DNA synthesis. Even small amounts of zinc are vital for maintaining a variety of biological processes(Bonaventura *et al*., 2015; Wessels *et al*., 2017). Other minerals such as potassium (K), magnesium (Mg), and iron (Fe) contribute additional nutritional benefits. Potassium is key in regulating blood pressure and heart function, magnesium supports enzymatic reactions and muscle health, and iron is critical for hemoglobin production and oxygen transport in the blood (Jomova *et al*., 2022).

### Amino acid profile

The result of the amino acid profile of the *M. oleifera* seed is presented in Table 5. The result showed that the most abundant essential amino acids is arginine while tryptophan is the least amino acid in the kernel and its defatted form. The defatting process significantly (p < 0.05) increase the overall essential amino acids. The highest arginine content was observed in the sample L, kernel defatted with mixture of solvents followed by Sample J, kernel defatted with n-hexane though not statistically significant (p > 0.05), while the lowest was in the undefatted seed, Sample I. Arginine is known to promote wound healing and immune function, which are crucial in combating the negative impacts of PEM (Stechmiller, 2010). The increase in arginine after defatting suggests enhanced bioavailability of this essential amino acid.

**Table 5:** Amino acids profile of *M. oleifera* kernel and the defatted kernels.

|  | Sample |  |  |  |
| --- | --- | --- | --- | --- |
|  | I | J | K | L |
| **Essential Amino Acids (EAA) g/100g protein** |  |  |  |  |
| Arginine | 6.76±0.02<sup>a</sup> | 7.44±0.03<sup>b</sup> | 7.18±0.02<sup>c</sup> | 7.66±0.03<sup>b</sup> |
| Histidine | 2.63±0.01<sup>a</sup> | 3.07±0.02<sup>b</sup> | 2.94±0.02<sup>b</sup> | 3.07±0.02<sup>b</sup> |
| Isoleucine | 3.20±0.02<sup>a</sup> | 3.72±0.01<sup>b</sup> | 3.42±0.01<sup>a</sup> | 3.72±0.01<sup>c</sup> |
| Leucine | 6.39±0.02<sup>a</sup> | 7.25±0.03<sup>b</sup> | 6.61±0.06<sup>a</sup> | 7.84±0.03<sup>c</sup> |
| Lysine | 3.75±0.01<sup>a</sup> | 4.06±0.02<sup>b</sup> | 3.88±0.04<sup>ab</sup> | 4.27±0.02<sup>c</sup> |
| Methionine | 1.22±0.00<sup>a</sup> | 1.29±0.01<sup>a</sup> | 1.27±0.01<sup>a</sup> | 1.33±0.02<sup>a</sup> |
| Phenylalanine | 4.21±0.02<sup>a</sup> | 4.85±0.01<sup>b</sup> | 4.74±0.02<sup>b</sup> | 5.03±0.04<sup>b</sup> |
| Threonine | 2.49±0.01<sup>a</sup> | 3.18±0.01<sup>b</sup> | 2.78±0.01<sup>c</sup> | 3.27±0.01<sup>b</sup> |
| Tryptophan | 0.74±0.00<sup>a</sup> | 0.85±0.01<sup>bc</sup> | 0.78±0.02<sup>ab</sup> | 0.95±0.01<sup>c</sup> |
| Valine | 4.08±0.01<sup>a</sup> | 4.28±0.01<sup>b</sup> | 4.40±0.01<sup>c</sup> | 4.70±0.01<sup>d</sup> |
| **Total of EAA (g/100g protein)** | **35.46±0.04<sup>a</sup>** | **39.97±0.05<sup>b</sup>** | **37.97±0.12<sup>c</sup>** | **41.81±0.15<sup>d</sup>** |
| **Non-Essential Amino Acids (NAA) g/100g protein** |  |  |  |  |
| Alanine | 3.44±0.01<sup>a</sup> | 4.07±0.03<sup>b</sup> | 3.63±0.02<sup>c</sup> | 4.17±0.02<sup>b</sup> |
| Aspartic acid | 8.19±0.02<sup>a</sup> | 9.18±0.04<sup>b</sup> | 8.59±0.02<sup>c</sup> | 9.47±0.04<sup>b</sup> |
| Cysteine | 0.60±0.00<sup>a</sup> | 0.94±0.02<sup>b</sup> | 0.82±0.02<sup>b</sup> | 1.20±0.00<sup>c</sup> |
| Glutamic acid | 10.39±0.08<sup>a</sup> | 11.54±0.02<sup>b</sup> | 10.66±0.03<sup>a</sup> | 12.20±0.08<sup>c</sup> |
| Glycine | 3.40±0.01<sup>a</sup> | 3.57±0.01<sup>b</sup> | 3.64±0.02<sup>bc</sup> | 3.83±0.02<sup>c</sup> |
| Proline | 3.06±0.01<sup>a</sup> | 3.43±0.01<sup>b</sup> | 3.24±0.0<sup>c</sup> | 3.47±0.01<sup>b</sup> |
| Serine | 3.19±0.01<sup>a</sup> | 4.06±0.02<sup>b</sup> | 3.68±0.01<sup>c</sup> | 4.06±0.02<sup>b</sup> |
| Tyrosine | 3.25±0.01<sup>a</sup> | 3.51±0.04<sup>ab</sup> | 3.37±0.04<sup>ab</sup> | 3.59±0.01<sup>b</sup> |
| **Total of NAA (g/100g protein)** | **35.50±0.04<sup>a</sup>** | **40.29±0.08<sup>b</sup>** | **37.61±0.02<sup>c</sup>** | **41.87±0.10<sup>d</sup>** |
| **Total** | **70.95±0.08<sup>a</sup>** | **80.25±0.13<sup>b</sup>** | **75.58±0.10<sup>c</sup>** | **83.68±0.04<sup>d</sup>** |

Values are expressed as means of three (3) replicates ± SEM (Standard Error of Mean). Values with different superscripts across the row are statistical ( $p < 0.05$ ) significant. I= undefatted *Moringa oleifera* seed, J= *Moringa oleifera* seed defatted with n-hexane, K= *Moringa oleifera* seed defatted with acetone, L= *Moringa oleifera* seed defatted with a mixture of n-hexane and acetone in ratio 1:1

Defatted samples J and L showed a significant increase (p < 0.05) in Histidine content compared to Sample I (*p* < 0.05), with Sample L having the highest value compared to undefatted and acetone defatted variants. Histidine is vital for growth and tissue repair, making it essential for recovery in PEM conditions (Siva Kiran *et al*., 2015).

Leucine and isoleucine are branched-chain amino acids (BCAAs) that play a key role in muscle protein synthesis. Leucine concentration significantly increased in mixture of solvent and n-hexane defatted variants, compared to undefatted and acetone defatted samples. This is particularly beneficial in addressing muscle wasting in PEM cases, where muscle breakdown exceeds muscle synthesis (Gorissen & Phillips, 2019).

Lysine, an essential amino acid important for collagen formation and protein utilization, increased significantly(p < 0.05) in defatted samples, with mixture of solvents defatted Sample showing the highest value followed by n-hexane defatted sample. In malnourished individuals, lysine deficiency can impair growth and immune responses (Morales *et al*., 2023).

The defatted process also significantly improve the Valine concentration with mixture of solvents defatted sample Sample L also showed the highest Valine value followed by n-hexane defatted sample, making them strong sources for improving the protein quality of diets for PEM treatment (Wei *et al*., 2021).

In general, the total Essential Amino Acids (EAA) concentration increased with defatting, which was significantly higher than the undefatted sample (*p* < 0.05). This improvement suggests that defatting *Moringa oleifera* seeds can enhance the nutritional quality of the seeds by concentrating the EAAs. This is crucial for addressing PEM, as EAAs are critical for muscle protein synthesis and metabolic recovery in malnourished individuals (Hoffer, 2016).

Non-essential amino acids, though synthesized by the body, contribute significantly to protein metabolism and overall health, especially in recovery from malnutrition. The result of the amino acids profile revealed that, glutamic acid and Cysteine were the most abundant and least abundant non-essential amino acids respectively. The defatted process significantly improve the overall non-essential amino acids. Alanine and aspartic acid are involved in glucose metabolism and energy production, which are critical in providing fuel for malnourished individuals(Remesar & Alemany, 2020). Cysteine is important for antioxidant production, particularly glutathione, which helps combat oxidative stress—a condition prevalent in PEM (Raj Rai *et al*., 2021). Glutamic acid serves as a precursor for glutamine, a vital nutrient for intestinal health, and is essential in PEM recovery due to its role in maintaining gut integrity and promoting immune function (Aanandhi & John, 2017). Proline and glycine are elevated after defatting, enhances the structural protein content, crucial for rebuilding tissues in malnourished individuals (Albaugh *et al*., 2017).

Protein-energy malnutrition is characterized by a significant deficiency of both protein and energy in the diet, leading to muscle wasting, weakened immunity, and poor growth, particularly in children. The data from this study demonstrates that defatting *Moringa oleifera* seeds significantly enhances the protein quality by increasing the concentration of both essential and non-essential amino acids, making it a potent nutritional supplement for individuals suffering from PEM. The increase in branched-chain amino acids (leucine, isoleucine, Valine) is particularly beneficial for promoting muscle protein synthesis and preventing muscle wasting in PEM (Rajendram *et al*., 2015). The higher levels of glutamic acid and cysteine also suggest that defatted *Moringa oleifera* seeds can help restore immune function and combat oxidative stress, which is critical in malnourished individuals (Oladele *et al*., 2022).

### In-vitro protein digestibility

Protein digestibility is an important metric, especially in the context of protein-energy malnutrition (PEM), as it determines how well proteins are broken down and absorbed, which is crucial for combating malnutrition. Figure 2 show the result of the in-vitro protein digestibility The *In-invitro* protein digestibility (IVPD) of the undefatted *Moringa oleifera* seed (Sample I) was significantly lower than the defatted samples (*p* < 0.05). This lower digestibility may be attributed to the presence of higher fat content, which can interfere with protein breakdown and enzymatic action during digestion (Orlien *et al*., 2023). Undefatted seeds likely retain anti-nutritional factors, such as phytates and saponins, which are known to reduce protein digestibility by binding to proteins and digestive enzymes, limiting their functionality (Ohanenye *et al*., 2022). In malnourished populations, consuming foods with low protein digestibility exacerbates protein deficiencies, making it difficult to meet dietary protein needs (Schönfeldt & Hall, 2012).

**Figure 2:**
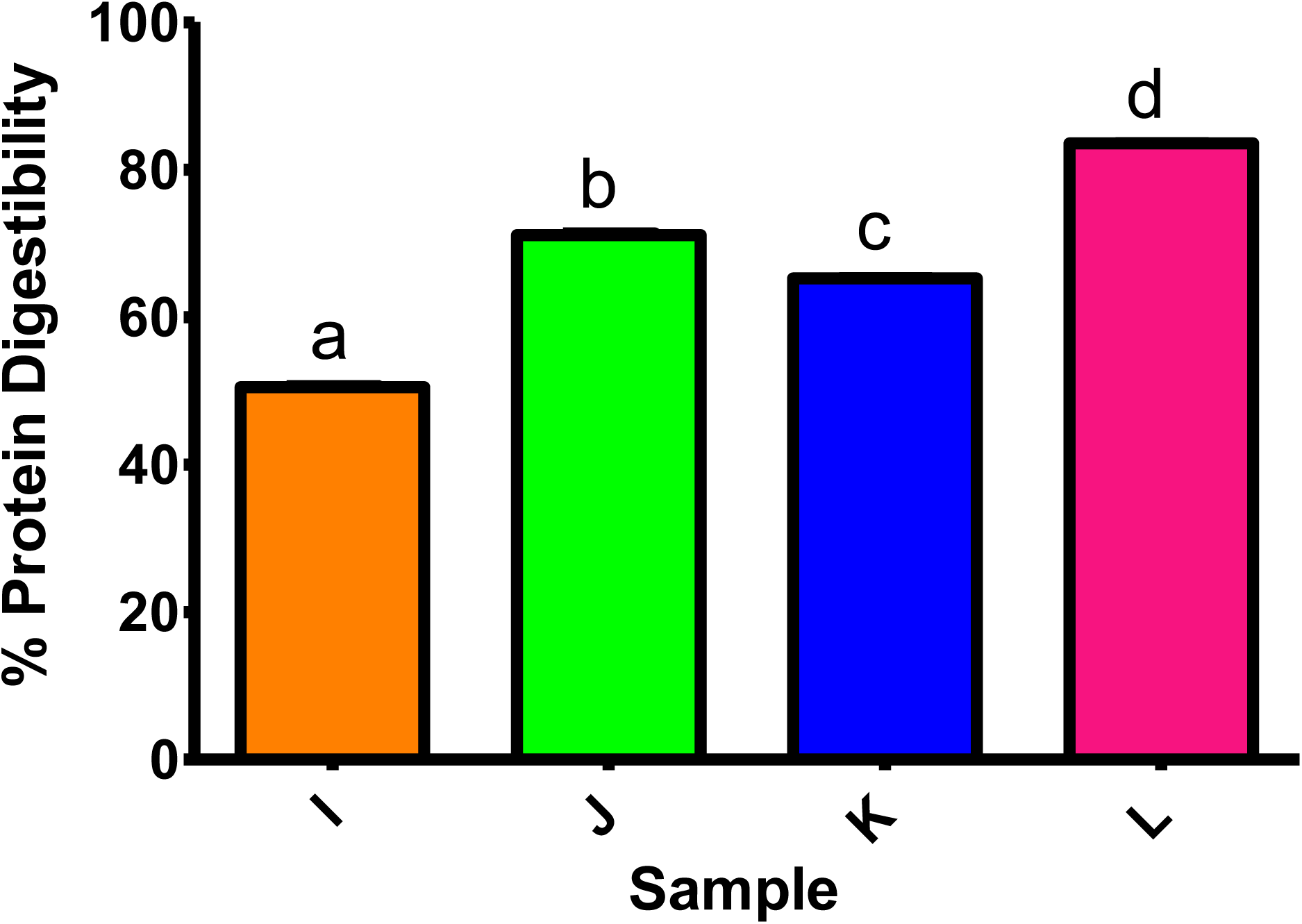
I*n-vitro* Protein Digestibility. Values are expressed as means of three (3) replicates ± SEM (Standard Error of Mean). Bars with different alphabets are statistically significant (p < 0.05) I= undefatted *Moringa oleifera* seed, J= *Moringa oleifera* seed defatted with n-hexane, K= *Moringa oleifera* seed defatted with acetone, L= *Moringa oleifera* seed defatted with a mixture of n-hexane and acetone in ratio 1:1.

The IVPD of seeds defatted with n-hexane (Sample J) was significantly higher indicating that defatting with n-hexane improves protein digestibility by removing fats that interfere with enzymatic action (*p* < 0.05). The removal of fats through solvent extraction enhances the access of proteolytic enzymes to protein molecules (Dzuvor *et al*., 2022). N-Hexane has been shown to effectively extract lipids without significantly denaturing proteins, preserving their structure and improving digestibility. In the context of PEM, diets formulated with defatted seeds could provide more bioavailable protein, which is crucial for restoring nitrogen balance and promoting growth (Chandran *et al*., 2023).

Sample K, defatted with acetone, showed a significantly (p < 0.05) lower digestibility than Sample J (defatted with n-hexane), though still significantly higher than the undefatted sample (*p* < 0.05). Acetone is a polar solvent and may have partially denatured some proteins during extraction, reducing their digestibility compared to n-hexane (Rose, 2019). Despite this, acetone extraction still improves digestibility by reducing the fat content. In terms of dietary application for protein-energy malnutrition, this sample would still be beneficial, though slightly less effective compared to n-hexane-defatted seeds in enhancing protein utilization (Ahmed *et al*., 2022; Elsebaie *et al*., 2023).

The highest protein digestibility was observed in Sample L, where a mixture of n-hexane and acetone (1:1 ratio) was used for defatting (*p* < 0.05). The combined use of these solvents likely maximized lipid removal without significant protein denaturation, providing the most accessible protein for digestion (Dzuvor *et al*., 2022).

Defatted Moringa seeds provide a lower fat content, allowing the digestive system to focus on protein breakdown. Additionally, removing fats can reduce the inhibitory effects of anti-nutritional factors on protein digestibility (Sardabi *et al*., 2022). For malnourished populations, consuming defatted Moringa seeds processed with optimal solvent extraction techniques could enhance protein utilization and improve nutritional outcomes(Gopalakrishnan *et al*., 2016).

### The bio-active compounds

The GC-MS analysis of *Moringa oleifera* seeds reveals a diverse range of bioactive compounds with potential pharmacological and industrial applications. This detailed chemical profile provides insight into the multifunctionality of the seeds in food, pharmaceutical, and cosmetic industries (Table 6).

**Table 6a:**
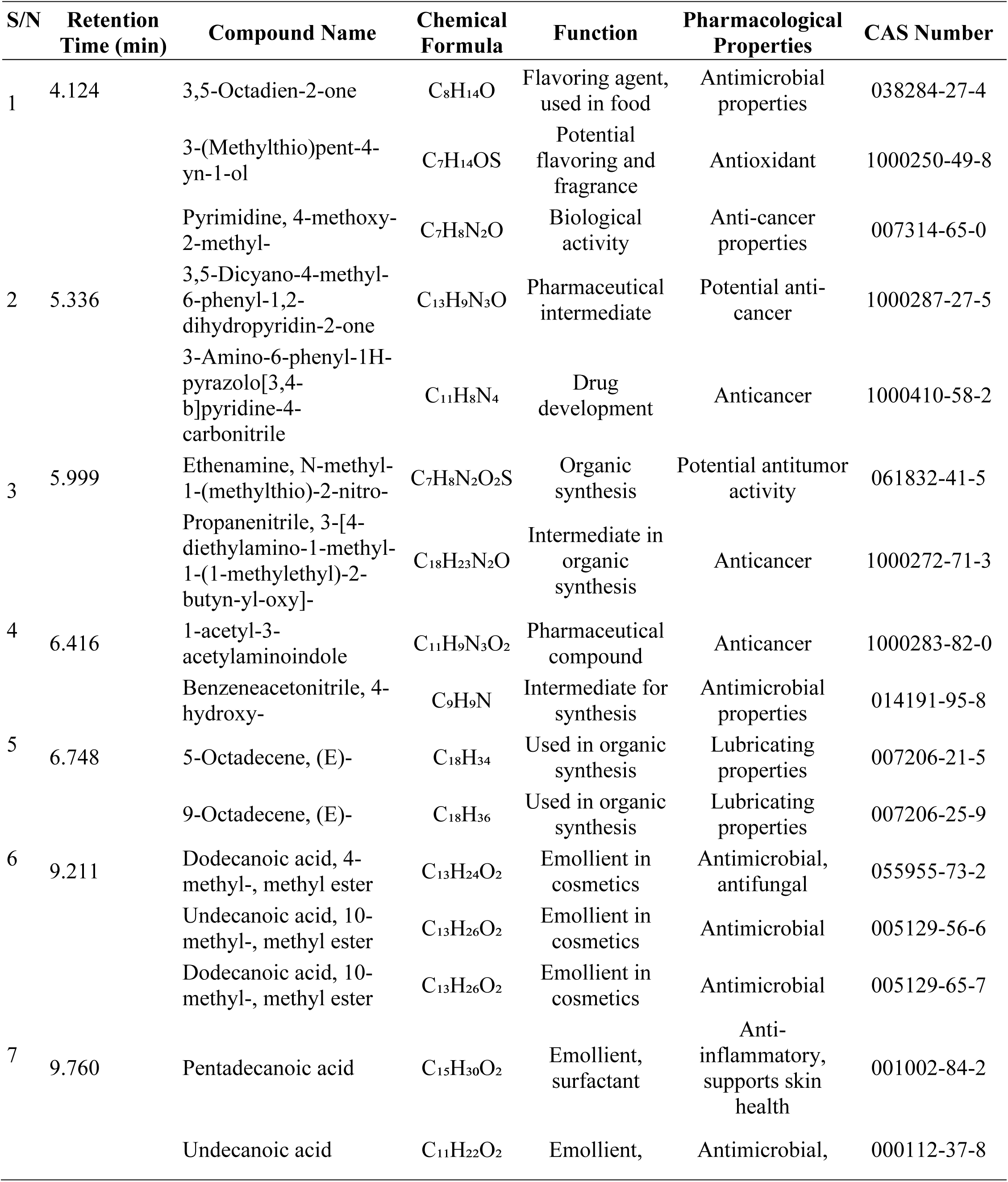
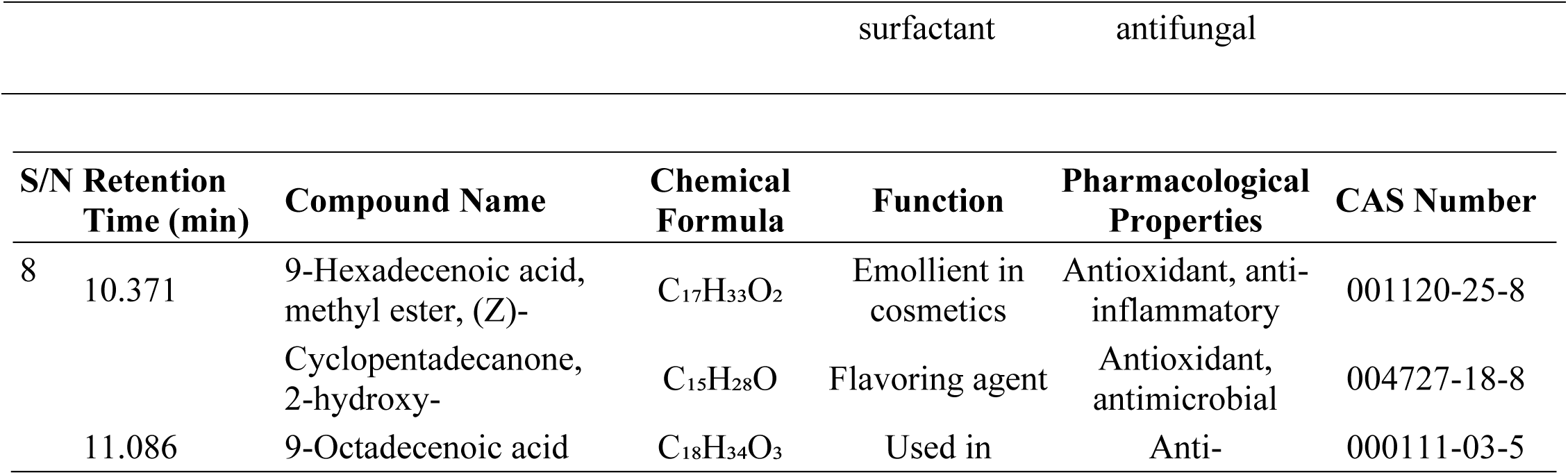
Bioactive compounds obtained from GC-MS analysis of n-hexane extract of *Moringa oleifera* kernel.

**Table 6b:** Bioactive compounds obtained from GC-MS analysis of n-hexane extract of *Moringa oleifera* kernel.

|  |  |  |  |  |  |  |
| --- | --- | --- | --- | --- | --- | --- |
| 9 |  | (Z)-, 2,3-dihydroxypropyl ester |  | cosmetics and foods | inflammatory, antioxidant |  |
| 10 | 12.069 | 1-(4-Amino-furazan-3-yl)-5-methyl-1H-[1,2,3]triazole-4-carboxylic acid amide | C<sub>9</sub>H<sub>8</sub>N<sub>4</sub>O<sub>2</sub> | Pharmaceutical compound | Antimicrobial | 1000300-80-6 |
|  |  | Thiophene-2-carboxylic acid, 4-bromo-3-methoxy- | C<sub>9</sub>H<sub>8</sub>BrN<sub>3</sub>O<sub>2</sub>S | Pharmaceutical compound | Anti-inflammatory | 110545-68-1 |
| 11 | 12.869 | Paromomycin | C<sub>21</sub>H<sub>43</sub>N<sub>7</sub>O<sub>13</sub> | Antibiotic | Antimicrobial | 007542-37-2 |
|  |  | 4-Hexenoic acid, 6-hydroxy-4-methyl-, methyl ester, (E)- | C<sub>9</sub>H<sub>16</sub>O<sub>3</sub> | Pharmaceutical compound | Anti-inflammatory | 053585-95-8 |
| 12 | 13.863 | 8-Methyl-6-nonenoic acid | C<sub>11</sub>H<sub>18</sub>O<sub>2</sub> | Flavoring agent | Antioxidant properties | 021382-25-2 |
|  |  | R(-)3,7-Dimethyl-1,6-octadiene | C<sub>18</sub>H<sub>34</sub> | Organic synthesis | Anti-inflammatory | 010281-56-8 |
| 13 | 15.395 | Succinic acid, 2,2,3,3-tetrafluoropropyl non-3-en-1-yl ester | C<sub>11</sub>H<sub>13</sub>F<sub>4</sub>O<sub>2</sub> | Chemical synthesis | Antimicrobial | 1000391-09-1 |
|  |  | Ethenamine, N-methyl-1-(methylthio)-2-nitro- | C<sub>7</sub>H<sub>8</sub>N<sub>2</sub>O<sub>2</sub>S | Intermediate for synthesis | Anticancer | 061832-41-5 |

### Flavoring Agents and Fragrance Compounds

The analysis identified several compounds with flavoring properties, such as 3,5-Octadien-2-one (C₈H₁₄O), which is commonly used in food as a flavoring agent and has known antimicrobial properties (CAS No. 038284-27-4). This aligns with previous studies that highlight the importance of volatile compounds in *Moringa oleifera* for their flavor-enhancing roles in culinary applications (Shunmugapriya *et al*., 2017). Similarly, Cyclopentadecanone, 2-hydroxy-(C₁₅H₂₈O), another flavoring agent, is noted for its antioxidant and antimicrobial properties (CAS No. 004727-18-8). These dual functionalities make these compounds valuable for both sensory enhancement and food preservation, where antimicrobial properties help in shelf-life extension (Shunmugapriya *et al*., 2017).

### Pharmaceutical Intermediates and Anticancer Potential

Several compounds identified have significant pharmaceutical relevance. For instance, 3,5-Dicyano-4-methyl-6-phenyl-1,2-dihydropyridin-2-one (C₁₃H₉N₃O) is noted for its role as a pharmaceutical intermediate with potential anti-cancer properties (CAS No. 1000287-27-5). Other compounds, such as 3-Amino-6-phenyl-1H-pyrazolo[3,4-b]pyridine-4-carbonitrile (C₁₁H₈N₄), have also demonstrated anticancer activities, reinforcing the potential of *Moringa oleifera* in drug discovery and development (CAS No. 1000410-58-2). These findings are consistent with the literature suggesting that *Moringa oleifera* seeds contain various bioactive compounds that exhibit antitumor effects (Tiloke *et al*., 2018).

### Antimicrobial and Antioxidant Properties

The seeds also contain compounds with antimicrobial and antioxidant properties. For instance, Pyrimidine, 4-methoxy-2-methyl-(C₇H₈N₂O) has anti-cancer properties (CAS No. 007314-65-0), while compounds such as Dodecanoic acid, 10-methyl-, methyl ester (C₁₃H₂₆O₂) show antimicrobial effects (CAS No. 005129-65-7). These compounds can be valuable in both pharmaceutical and cosmetic formulations, where their antimicrobial activities can enhance product safety and stability (Segwatibe *et al*., 2023). Additionally, the antioxidant properties of 9-Hexadecenoic acid, methyl ester, (Z)-(C₁₇H₃₃O₂) (CAS No. 001120-25-8) could contribute to the seeds’ protective effects against oxidative stress, a key factor in aging and disease development.

### Emollients and Surfactants in Cosmetics

Several fatty acids and their derivatives were identified as emollients and surfactants, including Pentadecanoic acid (C₁₅H₃₀O₂) and Undecanoic acid (C₁₁H₂₂O₂). Both are used in cosmetic formulations for their ability to soften and hydrate the skin (CAS No. 000112-37-8). These fatty acids also possess anti-inflammatory properties, which can help in the treatment of skin conditions such as eczema and dermatitis (Valarmathi *et al*., 2024).

### Organic Synthesis and Industrial Applications

Certain compounds, such as Ethenamine, N-methyl-1-(methylthio)-2-nitro-(C₇H₈N₂O₂S) (CAS No. 061832-41-5), were highlighted for their roles in organic synthesis. This compound, in particular, shows potential antitumor activity, which could make it a valuable intermediate in drug synthesis. Similarly, other long-chain hydrocarbons like 5-Octadecene (C₁₈H₃₄) (CAS No. 007206-21-5) and 9-Octadecene (C₁₈H₃₆) (CAS No. 007206-25-9) are known for their lubricating properties, which could find applications in industrial formulations and biolubricants (Gharsallah *et al*., 2023).

The GC-MS analysis of *Moringa oleifera* seeds demonstrates a rich chemical profile with a wide range of applications.

### Body Weight

The figure 3 shows the effects of soya bean and defatted *Moringa oleifera* seed-based diets, on the weekly mean body weight of protein-energy malnourished rats over a four-week period. As shown, the control group, which was fed a commercial diet, experienced a steady and significant weight gain. This is expected, as the control group had access to a nutritionally balanced diet that supported normal growth and weight gain in the rats (Sousa *et al*., 2018).

**Figure 3:**
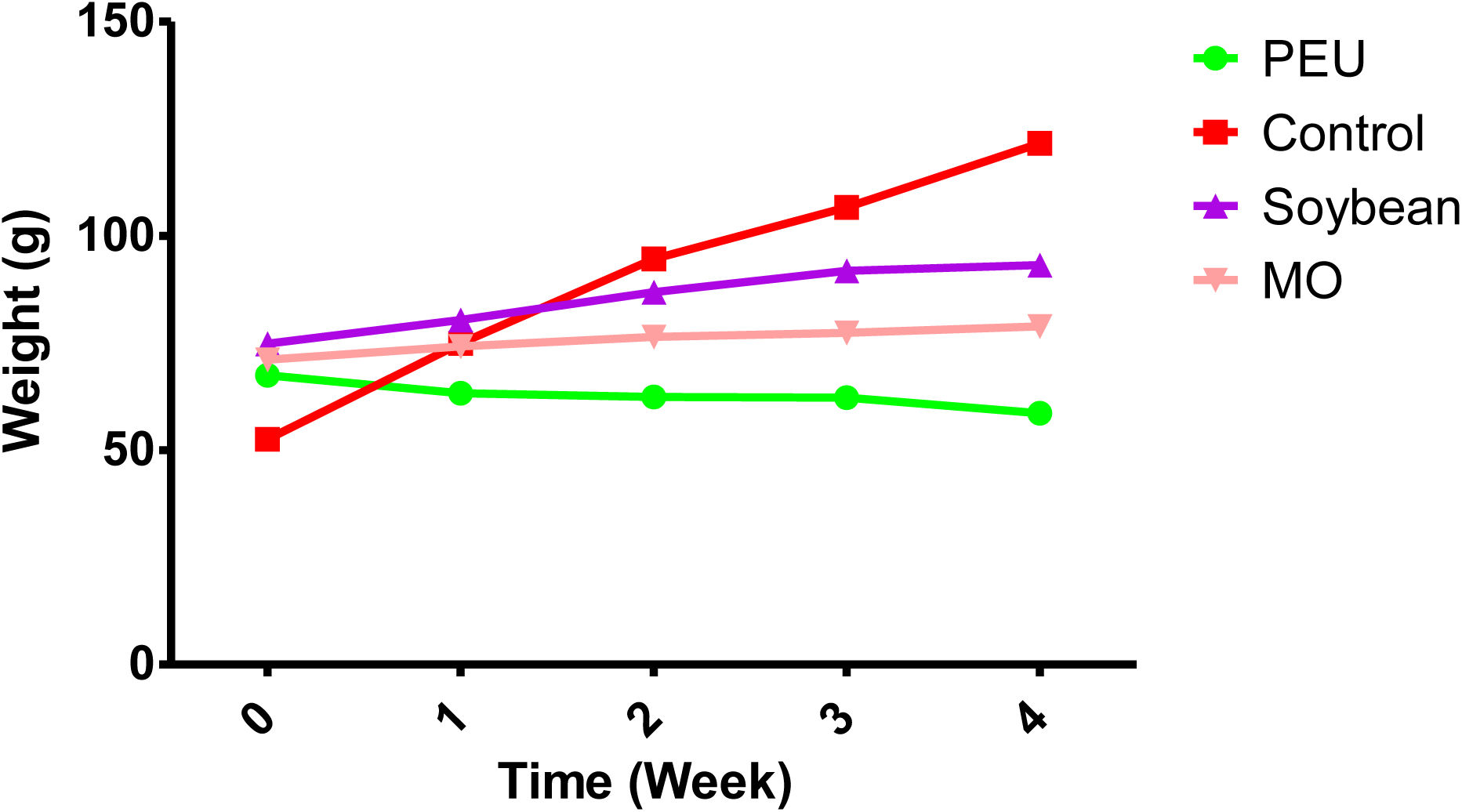
Effects of *M. oleifera* kernel based-diet on mean weekly body weight of protein energy malnourished rats. Values are expressed as means of five (5) replicates ± SEM (Standard Error of Mean). Control= Control Animals fed with commercial diet, PEU= Animals malnourished with low protein isocaloric based diet, Soya bean = undernourished animals treated with 15% soya been, MO= undernourished animals treated with 15% defatted *Moringa oleifera* seed.

In contrast, the PEU (Protein Energy Undernourished) group, which was fed a low-protein isocaloric diet, showed little to no weight gain, maintaining an almost flat line throughout the study. This result is typical of protein-energy malnutrition, where a lack of adequate protein leads to stunted growth and muscle wasting. This supports the understanding that an insufficient protein supply directly affects weight maintenance in animals(de França *et al*., 2009).

The groups treated with 15% soybean and 15% defatted *Moringa oleifera* seed (MO) showed significant (p < 0.05) increases in body weight compared to protein energy undernourished group. Previous studies indicate that soybeans and *Moringa oleifera* seeds are rich in high-quality protein and essential amino acids, which promote muscle mass and overall growth(Abdel-Latif *et al*., 2022).

However, *Moringa oleifera* seeds, though rich in nutrients, may have some anti-nutritional factors that reduce their efficacy compared to soybean, as noted in other studies. The partial defatting of *Moringa oleifera* may improve its protein content but could also reduce some lipid-based nutrients necessary for optimal growth(Soetan & Aiyelaagbe, 2016).

### Biochemical Analyses Liver Function

Table 7 shows the results of the effects of defatted *Moringa oleifera* kernel based-diet on liver function parameters of protein energy malnourished rats. The parameters measured—AST, ALT, ALP, TB, CB, TP, ALB, and GLO—are key indicators of liver health and metabolic function. The results show significant variations across the experimental groups, indicating both the effects of malnutrition and the efficacy of the treatments administered.

**Table 7:** Effects of *Moringa oleifera* kernel based-diet on liver function indices of protein energy malnourished rats.

| Group | AST (U/L) | ALT (U/L) | ALP (U/L) | TB (mg/dL) | CB (mg/dL) | TP (g/dL) | ALB (g/dL) | GLO (g/dL) |
| --- | --- | --- | --- | --- | --- | --- | --- | --- |
| A | 249.75±7.05<sup>a</sup> | 105.50±9.15<sup>a</sup> | 254.50±13.01<sup>a</sup> | 6.48±1.44<sup>a</sup> | 2.90±0.80<sup>a</sup> | 71.00±4.80<sup>a</sup> | 37.75±1.80<sup>a</sup> | 33.25±3.07<sup>a</sup> |
| B | 296.75±20.70<sup>b</sup> | 121.00±14.34<sup>b</sup> | 741.25±40.44<sup>b</sup> | 15.60±3.15<sup>b</sup> | 6.20±1.28<sup>b</sup> | 60.00±1.08<sup>b</sup> | 31.50±0.65<sup>b</sup> | 28.50±0.50<sup>b</sup> |
| C | 267.00±13.89<sup>ab</sup> | 106.25±12.32<sup>a</sup> | 491.50±7.60<sup>b</sup> | 13.80±1.21<sup>b</sup> | 5.93±0.27<sup>b</sup> | 73.25±2.87<sup>a</sup> | 36.50±1.04<sup>a</sup> | 36.75±2.02<sup>a</sup> |
| D | 213.75±2.63<sup>c</sup> | 92.75±3.15<sup>c</sup> | 318.25±9.00<sup>ac</sup> | 6.15±0.46<sup>a</sup> | 3.08±0.287<sup>a</sup> | 77.00±2.08<sup>a</sup> | 39.75±1.11<sup>a</sup> | 37.25±1.18<sup>a</sup> |

Values are expressed as means of five (5) replicates ± SEM (Standard Error of Mean). Values with different superscripts down the column are statistical (p < 0.05) significant. AST= aspartate amino transferase, ALT= alanine amino transferase, ALP = Alkaline Phosphatase, TB = total bilirubin CB= conjugated bilirubin, TP= total protein, ALB= albumin, GLO= globulin: A= Control Animals fed with commercial diet, B= Animals malnourished with low protein isocaloric based diet, C= undernourished animals treated with 15% soya been, D= undernourished animals treated with 15% defatted *Moringa oleifera* seed

**Table 8:** Effects of *Moringa oleifera* kernel based-diet on Kidney function indices of protein energy malnourished rats.

| Group | Na (mEq/L) | K (mEq/L) | Cl<sup>-</sup> (mEq/L) | HCO<sub>3</sub><sup>-</sup> (mEq/L) | Ca<sup>2+</sup> (mg/dL) | Urea (mg/dL) | Creatinine (μmol/L) |
| --- | --- | --- | --- | --- | --- | --- | --- |
| A | 122.00±0.41<sup>a</sup> | 6.20±0.04<sup>b</sup> | 95.75±0.48<sup>a</sup> | 21.00±0.41<sup>a</sup> | 1.17±0.01<sup>a</sup> | 8.15±1.07<sup>a</sup> | 72.25±4.07<sup>a</sup> |
| B | 106.00±0.41<sup>b</sup> | 5.70±0.04<sup>c</sup> | 79.50±0.65<sup>b</sup> | 19.75±0.25<sup>b</sup> | 0.99±0.03<sup>b</sup> | 9.80±0.54<sup>a</sup> | 113.00±2.12<sup>b</sup> |
| C | 137.75±0.63<sup>c</sup> | 8.83±0.09<sup>a</sup> | 101.50±0.65<sup>a</sup> | 26.25±0.85<sup>c</sup> | 1.17±0.01<sup>a</sup> | 8.40±0.52<sup>a</sup> | 90.50±4.87<sup>a</sup> |
| D | 141.50±1.32<sup>c</sup> | 9.08±0.30<sup>a</sup> | 98.50±0.65<sup>a</sup> | 26.25±1.11<sup>c</sup> | 1.20±0.02<sup>a</sup> | 5.10±0.44<sup>b</sup> | 52.50±4.83<sup>c</sup> |

Values are expressed as means of five (5) replicates ± SEM (Standard Error of Mean). Values with different superscripts down the column are statistical (p < 0.05) significant: A= Control Animals fed with commercial diet, B= Animals malnourished with low protein isocaloric based diet, C= undernourished animals treated with 15% soya been, D= undernourished animals treated with 15% defatted *Moringa oleifera* seed

AST and ALT are liver enzymes that serve as markers of hepatocellular injury. Elevated levels typically indicate liver damage or inflammation. In this study, group B (malnourished rats fed with a low-protein isocaloric diet) had significantly (p < 0.05) elevated AST and ALT levels compared to the control (group A), reflecting hepatic stress induced by protein-energy malnutrition (PEM).

The rats treated with 15% soybean (group C) exhibited moderate AST and ALT levels compared to group B, suggesting partial recovery of liver function. However, the *Moringa oleifera* kernel based diet-treated group (D) demonstrated the most significant (p < 0.05) reduction in AST and ALT, approaching normal levels. This indicates the potential hepatoprotective effects of *Moringa oleifera*, possibly due to its antioxidant and anti-inflammatory properties, which have been reported in previous studies(Atta *et al*., 2017).

ALP is another marker of liver function, often elevated in conditions involving bile duct obstruction or hepatobiliary disease. In group B, malnourished group, ALP levels were significantly (p < 0.05) elevated, indicating severe hepatobiliary dysfunction due to malnutrition. However, treatment with both soybean based-diet (group C) and defatted *Moringa oleifera* kernel based-diet (group D) significantly (p < 0.05) lowered ALP levels, with group D showing the greatest improvement. The reduction in ALP suggests that both diets improved bile flow and overall liver function, with *Moringa oleifera* being more effective in restoring liver enzyme levels to near-normal values(Ekong *et al*., 2022).

Total bilirubin (TB) and conjugated bilirubin (CB) are markers of liver excretory function. Elevated levels of bilirubin, as observed in group B (malnourished group), indicate impaired liver function and bile excretion. These elevated bilirubin levels confirm the presence of cholestasis or hepatic damage due to malnutrition. In contrast, groups C and D showed significant (p < 0.05) reductions in both total and conjugated bilirubin levels. The values of Group D are not significant (p > 0.05) different compared to the control values, further indicating that *Moringa oleifera* seed has a restorative effect on liver excretory function (Mhlomi *et al*., 2022).

Total protein, albumin, and globulin levels are crucial indicators of nutritional status and liver synthetic function. Protein-energy malnutrition leads to a decline in protein synthesis, reflected in the significant (p < 0.05) reductions in Total Protein, Albumin, and Globulin in group B (malnourished rats). The decrease in these proteins compromises immune function and fluid balance, as albumin helps maintain osmotic pressure(Amir, 2003). Rats in group C (treated with 15% soybean) and group D (treated with Moringa oleifera) showed improved protein levels. Group D displayed higher Total protein, albumin and globulin compared to group C, suggesting that *Moringa oleifera* was more effective in promoting protein synthesis and liver function restoration. This could be due to the high nutritional content and bioactive compounds found in *Moringa oleifera*, including essential amino acids and phytochemicals that support protein metabolism (Ayodele *et al*., 2021).

### Kidney function

The data in figure 8 shows the kidney function analysis in protein-energy undernourished rats treated with soybean-based diets (Group C) and Moringa oleifera-based diets (Group D). These groups are compared with control animals (Group A) and malnourished animals (Group B). The analysis of key biochemical markers including sodium (Na), potassium (K), chloride (Cl-), bicarbonate (HCO₃⁻), calcium (Ca²⁺), urea, and creatinine reveals significant alterations in kidney function due to undernutrition and dietary intervention.

Group B (malnourished) exhibited a significantly (p < 0.05) reduced concentration of all the electrolytes (Na^+^, K^+^, Ca^2+^,Cl^-^, and HCO_3_^-^) studied compared to the control group A, indicating an imbalance, impaired acid-base homeostasis and renal function due to protein-energy malnutrition. This aligns with previous studies showing that malnutrition leads to disturbances in electrolyte balance (Khan *et al*., 2021), and suggests that PEM affects potassium retention, leading to hypokalemia, which could compromise muscle and heart function. Treatment with soya bean and defatted *M. oleifera* based-diets as seen in Both Group C and Group D showed significantly (p < 0.05) elevated of these electrolytes levels compared to Group A, suggesting that dietary intervention with 15% soybean and defatted *Moringa oleifera* seed could improve electrolytes homeostasis. However, the higher values in Group D suggest that *Moringa oleifera* may be more effective in restoring there levels(Abdel-Latif *et al*., 2022).

The kidney function result also revealed significant (p < 0.05) elevated urea and creatinine levels in Group B compared to Group A, suggesting impaired renal clearance, reduced kidney filtration capacity and compromised glomerular function due to malnutrition. Urea, a byproduct of protein metabolism, is elevated when kidney function is compromised (Adeyomoye *et al*., 2022). In contrast, Group D showed a significant reduction in urea levels compared to the control group (*p* < 0.05). On the other hand, Both Group C and D showed improved creatinine levels with Group D showing a significant reduction compared to the control (*p* < 0.05). This suggests that *Moringa oleifera*-based diets may offer greater protection against kidney damage than soybean diets, possibly due to its antioxidant and anti-inflammatory properties (El Sohaimy *et al*., 2015). Suggesting that *Moringa oleifera* improves kidney function by enhancing urea and creatinine excretion, potentially due to its nephroprotective properties (Akinrinde *et al*., 2020).

Protein-energy malnutrition significantly impairs kidney function, as evidenced by altered serum electrolyte levels, increased urea, and creatinine. However, dietary interventions using *Moringa oleifera* and soybean-based diets effectively improved most of the kidney function markers, with *Moringa oleifera* showing superior nephroprotective effects. These findings support the potential therapeutic use of *Moringa oleifera* for ameliorating renal dysfunction in PEM cases(Akinrinde *et al*., 2020).

### Plasma lipid profile

Table 9 shows the effects of defatted *Moringa oleifera* seed-based diets on the lipid profile of protein-energy malnourished rats, with parameters; Total Cholesterol (TC), Triglycerides (TG), High-Density Lipoprotein (HDL), Low-Density Lipoprotein (LDL), the Atherogenic Index of Plasma (AIP), and the Atherogenic Coefficient (AC).

**Table 9:** Effects of *Moringa oleifera* kernel based-diet on lipid profile of protein energy malnourished rats.

| Group | TC (mg/dL) | TG (mg/dL) | HDL (mg/dL) | LDL (mg/dL) | AIP | AC |
| --- | --- | --- | --- | --- | --- | --- |
| A | 122.00 ± 2.16<sup>a</sup> | 100.75 ± 4.15<sup>a</sup> | 47.25 ± 2.72<sup>a</sup> | 66.75 ± 4.55<sup>a</sup> | 0.33 ± 0.05<sup>a</sup> | 1.58 ± 0.1<sup>a</sup> |
| B | 135.75 ± 2.56<sup>c</sup> | 105.75 ± 4.46<sup>a</sup> | 31.75 ± 0.85<sup>b</sup> | 81.33 ± 3.97<sup>b</sup> | 0.52 ± 0.06<sup>b</sup> | 3.28 ± 0.12<sup>b</sup> |
| C | 111.00 ± 3.63<sup>d</sup> | 82.50 ± 1.04<sup>b</sup> | 38.25 ± 1.80<sup>b</sup> | 65.85 ± 1.84<sup>a</sup> | 0.33 ± 0.04<sup>a</sup> | 1.90 ± 0.09<sup>a</sup> |
| D | 100.00 ± 0.91<sup>b</sup> | 75.50 ± 2.96<sup>b</sup> | 48.00 ± 1.08<sup>a</sup> | 56.08 ± 0.48<sup>a</sup> | 0.20 ± 0.03<sup>b</sup> | 1.08 ± 0.08<sup>a</sup> |

Values are expressed as means of five (5) replicates ± SEM (Standard Error of Mean). Values with different superscripts down the column are statistical (p < 0.05) significant. HDL= High Density Lipoprotein, TG= Triglyceride, TC= Total Cholesterol, LDL = Low Density Lipoprotein, AIP= Atherogenic Index of Plasma, AC= Atherogenic Coefficient: A= Control Animals fed with commercial diet, B= Animals malnourished with low protein isocaloric based diet, C= undernourished animals treated with 15% soya been, D= undernourished animals treated with 15% defatted *Moringa oleifera* seed

The rats malnourished with a low-protein isocaloric diet (Group B), has significantly (p < 0.05) elevated Total Cholesterol, Triglyceride and Low Density Lipoprotein-cholesterol with significantly (p < 0.05) reduced High Density Lipoprotein-cholesterol levels compared to the control group A. The Atherogenic Index of Plasma (AIP) and Atherogenic Coefficient (AC) are significantly (p < 0.05) increased in malnourished group B compared to the control (group A). These values indicate a heightened risk of cardiovascular disease. This suggests that protein-energy malnutrition induces dyslipidemia, impairing lipid metabolism and leading to elevated atherogenic risk (Ofori, 2023).

Group C, malnourished rats treated with a 15% soybean-based diet, showed significant improvement in their lipid profile. The Total Cholesterol and Triglyceride levels were significantly (p < 0.05) lower than Group B. HDL significantly (p < 0.05) increased with a significant (p < 0.05) decrease in LDL as compared to the malnourished group (B). The Atherogenic Index of Plasma and Atherogenic Coefficient significantly (p < 0.05) decreased, indicating improved cardiovascular health compared to the malnourished group, highlighting the lipid-lowering and cardioprotective effects of soybean in the diet (Ofori, 2023).

Group D, malnourished rats treated with a 15% defatted *Moringa oleifera* seed-based diet, had the most favorable lipid profile among the treatment groups. TC, TG, and LDL levels were significantly (p < 0.05) lowered. HDL significantly (p < 0.05) increase as compared to all other groups. The AIP and AC values significantly (p < 0.05) decrease, indicating a substantial reduction in cardiovascular risk. This improvement may be attributed to the high antioxidant constituents in Moringa oleifera, which enhances lipid metabolism and reduces oxidative stress, leading to improved cardiovascular outcomes in malnourished rats(Asogwa *et al*., 2017).

### Conclusion

*Moringa oleifera* seeds, when defatted, present a cost-effective and nutritionally rich option for the management of protein energy malnutrition. Given the accessibility and affordability of *Moringa oleifera* in regions prone to malnutrition, its incorporation into dietary interventions could have significant impacts on reducing the prevalence of PEM and associated health complications. Further research and clinical trials are recommended to confirm its efficacy in human populations and explore its broader applications in nutritional therapies.

## References

Aanandhi, M. V., & John, M. R. (2017). Role of Glutamine Supplementation in Critically ill patients: A Narrative Review. Research Journal of Pharmacy and Technology, 10(8), 2801–2808.

Abdel-Latif, H. M. R., Abdel-Daim, M. M., Shukry, M., Nowosad, J., & Kucharczyk, D. (2022). Benefits and applications of Moringa oleifera as a plant protein source in Aquafeed: A review. Aquaculture, 547, 737369.

Adelusi, S. A., & Olowokere, J. O. (1985). A rapid method of induction of post-natal protein-energy malnutrition (PEM) in laboratory animals. Niger J Appl Sci, 3, 171–174.

Adeniyi, S. A., Orjiekwe, C. L., Ehiagbonare, J. E., & Arimah, B. D. (2012). Evaluation of chemical composition of the leaves of Ocimum gratissimum and Vernonia amygdalina. International Journal of Biological and Chemical Sciences, 6(3), 1316–1323.

Adeyomoye, O. I., Akintayo, C. O., Omotuyi, K. P., & Adewumi, A. N. (2022). The biological roles of urea: A review of preclinical studies. Indian Journal of Nephrology, 32(6), 539–545.

Adibah, K. Z. M., & Azzreena, M. A. (2019). Plant toxins: alkaloids and their toxicities. GSC Biological and Pharmaceutical Sciences, 6(2).

Ahlers, J. (1975). The mechanism of hydrolysis of β-glycerophosphate by kidney alkaline phosphatase. Biochemical Journal, 149(3), 535–546.

Ahmed, M. H., Vasas, D., Hassan, A., & Molnár, J. (2022). The impact of functional food in prevention of malnutrition. PharmaNutrition, 19, 100288.

Akinrinde, A. S., Oduwole, O., Akinrinmade, F. J., & Bolaji-Alabi, F. B. (2020). Nephroprotective effect of methanol extract of Moringa oleifera leaves on acute kidney injury induced by ischemia-reperfusion in rats. African Health Sciences, 20(3), 1382–1396.

Albaugh, V. L., Mukherjee, K., & Barbul, A. (2017). Proline precursors and collagen synthesis: biochemical challenges of nutrient supplementation and wound healing. The Journal of Nutrition, 147(11), 2011–2017.

Ali, A. A. H. (2023). Overview of the vital roles of macro minerals in the human body. Journal of Trace Elements and Minerals, 4, 100076.

Allain, C. C., Poon, L. S., Chan, C. S. G., Richmond, W., & Fu, P. C. (1974). Enzymatic determination of total serum cholesterol. Clinical Chemistry, 20(4), 470–475.

Amir, M. A. (2003). Effects Of Protein-Energy Malnutrition On The Immune System. Makara Journal of Science, 7(2), 4.

Anselm, O. U., Dixon, O. N., & Ugochi, U. (2023). Nutrient assessment of gluten free biscuit from African pear and orange flesh sweet potato flour blends. J. Agric. Food Sci. Biotechnol, 1, 83–94.

Ashraf, F., & Gilani, S. R. (2007). Fatty acids in Moringa oleifera oil. Jour. Chem. Soc. Pak, 29.

Asogwa, I. S., Ani, J. C., & Okoye, E. C. (2017). The Hypolipidemic Effect of Moringa Leaf Powder Supplementation in High Fat Diet Fed Rats. Journal of Nutritional Ecology and Food Research, 4(2), 161–166.

Atta, A. H., Soufy, H., Nasr, S. M., Soliman, A. M., Nassar, S. A., Maweri, A., Abd El-Aziz, T. H., Desouky, H. M., & Abdalla, A. M. (2017). Hepatoprotective and antioxidant effects of methanol extractof Moringa oleifera leaves in rats. Wulfenia, 24(3), 249–268.

Awuchi, C. G., Igwe, V. S., & Amagwula, I. O. (2020). Ready-to-use therapeutic foods (RUTFs) for remedying malnutrition and preventable nutritional diseases. International Journal of Advanced Academic Research, 6(1), 47–81.

Ayodele, P. F., Oyedotun, O., Onifade, O. F., Adeosun, A. M., Odeniyi, I. A., Omowaye, O. S., & Akram, M. (2021). Nutritional Evaluation of Moringa oleifera Leaves and the Effect of Its Bio-fortification with Animal Feed on Physical Changes and Organ Weights in Male Albino Rats. Asian Journal of Research in Biochemistry, 1–8. 10.9734/ajrb/2021/v9i430206

Bartels, H., Bohmer, M., & Heierli, C. (1972). Estimation of serum creatinine without removal of protein.

Boham, B. A., & Kocipai-Abyazan, R. (1974). Flavonoids and condensed tannins from leaves of Hawaiian vaccinium vaticulatum and V. calycinium. Pacific Sci, 48(4), 458–463.

Bonaventura, P., Benedetti, G., Albarède, F., & Miossec, P. (2015). Zinc and its role in immunity and inflammation. Autoimmunity Reviews, 14(4), 277–285.

Busch, K. W., Howell, N. G., & Morrison, G. H. (1974). Simultaneous determination of electrolytes in serum using a vidicon flame spectrometer. Analytical Chemistry, 46(9), 1231–1236.

Chandran, A. S., Suri, S., & Choudhary, P. (2023). Sustainable plant protein: an up-to-date overview of sources, extraction techniques and utilization. Sustainable Food Technology, 1(4), 466–483.

de França, S. A., Dos Santos, M. P., Garófalo, M. A. R., Navegantes, L. C., do Carmo Kettelhut, I., Lopes, C. F., & Kawashita, N. H. (2009). Low protein diet changes the energetic balance and sympathetic activity in brown adipose tissue of growing rats. Nutrition, 25>(11–12), 1186–1192.

Dhakar, R., Pooniya, B., Gupta, M., Maurya, S., Bairwa, N., & Sanwarmal. (2011). Moringa: The herbal gold to combat malnutrition. Chronicles of Young Scientists, 2(3), 119. 10.4103/2229-5186.90887

Djurdjević, L., Mitrović, M., & Pavlović, P. (2007). Total phenolics and phenolic acids in plants and soils. In Cell Diagnostics (pp. 155–168). CRC Press.

Doumas, B. T. (1975). Standards for total serum protein assays: a collaborative study. Clinical Chemistry, 21(8), 1159–1166. 10.1093/clinchem/21.8.1159

Dowd, M. K., Boykin, D. L., Meredith Jr, W. R., Campbell, B. T., Bourland, F. M., Gannaway, J. R., Glass, K. M., & Zhang, J. (2010). Fatty acid profiles of cottonseed genotypes from the national cotton variety trials.

Dzuvor, C. K. O., Pan, S., Amanze, C., Amuzu, P., Asakiya, C., & Kubi, F. (2022). Bioactive components from Moringa oleifera seeds: production, functionalities and applications–a critical review. Critical Reviews in Biotechnology, 42(2), 271–293.

Ekong, A. I., Bassey, E. E., Achor, A. B., & Francis, A. (2022). Effects of weaning diet s supplemented with Moringa oleifera leaf powder on the biochemical an d hematological indices of weanling Wistar rats. Sch. Int. J. Biochem, 5, 50–56.

El Sohaimy, S. A., Hamad, G. M., Mohamed, S. E., Amar, M. H., & Al-Hindi, R. R. (2015). Biochemical and functional properties of Moringa oleifera leaves and their potential as a functional food. Global Advanced Research Journal of Agricultural Science, 4(4), 188–199.

Elsebaie, E. M., Abdel-Fattah, A. N., Bakr, N. A., Attalah, K. M., & Aweas, A.-H. A. (2023). Principles of Nutritional Management in Patients with Liver Dysfunction—A Narrative Review. Livers, 3(2), 190–218.

Friedewald, W. T., Levy, R. I., & Fredrickson, D. S. (1972). Estimation of the concentration of low-density lipoprotein cholesterol in plasma, without use of the preparative ultracentrifuge. Clinical Chemistry, 18(6), 499–502.

Gharsallah, K., Rezig, L., Rajoka, M. S. R., Mehwish, H. M., Ali, M. A., & Chew, S. C. (2023). Moringa oleifera: Processing, phytochemical composition, and industrial application. South African Journal of Botany, 160, 180–193.

Goldenberg, H., & Drewes, P. A. (1971). Direct photometric determination of globulin in serum. Clinical Chemistry, 17(5), 358–362. 10.1093/clinchem/17.5.358

Gopalakrishnan, L., Doriya, K., & Kumar, D. S. (2016). Moringa oleifera: A review on nutritive importance and its medicinal application. In Food Science and Human Wellness (Vol. 5, Issue 2, pp. 49–56). Elsevier B.V. 10.1016/j.fshw.2016.04.001

Gorissen, S. H. M., & Phillips, S. M. (2019). Branched-chain amino acids (leucine, isoleucine, and valine) and skeletal muscle. In Nutrition and skeletal muscle (pp. 283–298). Elsevier.

Gornall, A. G., Bardawill, C. J., & David, M. M. (1949). Determination of serum proteins by means of the biuret reaction. J. Biol. Chem, 177(2), 751–766.

Grases, F., Prieto, R. M., & Costa-Bauza, A. (2017). Dietary phytate and interactions with mineral nutrients. Clinical Aspects of Natural and Added Phosphorus in Foods, 175–183.

Gulati, P., Li, A., Holding, D., Santra, D., Zhang, Y., & Rose, D. J. (2017). Heating reduces proso millet protein digestibility via formation of hydrophobic aggregates. Journal of Agricultural and Food Chemistry, 65(9), 1952–1959.

Harborne, J. B., & Mabry, T. J. (2013). The flavonoids: advances in research.

Harland, B. F., Oberleas, D., & J, C. E. R. G. J. G. D. P. K. R. G. S. B. G. S. B. T. K. D. Z. (1986). Anion-exchange method for determination of phytate in foods: collaborative study. Journal of the Association of Official Analytical Chemists, 69(4), 667–670.

Hoffer, L. J. (2016). Human protein and amino acid requirements. Journal of Parenteral and Enteral Nutrition, 40(4), 460–474.

Horwitz, W., & Latimer, G. (2005). AOAC International: Gaithersburg. MD, USA, 18.

Ibukun, E., Edobor, G., & Adeleke Ojo, O. (2013). Qualitative And Quantitative Analysis Of Phytochemicals In Senecio Biafrae Leaf International Journal Of Inventions In Pharmaceutical Sciences Qualitative And Quantitative Analysis Of Phytochemicals In Senecio Biafrae Leaf. Int.J.Inv.Pharm.Sci, 1(5), 428–432. www.ijips.net

Izzo, C., Grillo, F., & Murador, E. (1981). Improved method for determination of high-density-lipoprotein cholesterol I. Isolation of high-density lipoproteins by use of polyethylene glycol 6000. Clinical Chemistry, 27(3), 371–374.

Jomova, K., Makova, M., Alomar, S. Y., Alwasel, S. H., Nepovimova, E., Kuca, K., Rhodes, C. J., & Valko, M. (2022). Essential metals in health and disease. Chemico-Biological Interactions, 367, 110173.

Jørgensen, K., & Astrup, P. (1957). Standard bicarbonate, its clinical significance, and a new method for its determination. Scandinavian Journal of Clinical and Laboratory Investigation, 9(2), 122–132.

Kamran, M., Hussain, S., Abid, M. A., Syed, S. K., Suleman, M., Riaz, M., Iqbal, M., Mahmood, S., Saba, I., & Qadir, R. (2020). Phytochemical composition of moringa oleifera its nutritional and pharmacological importance. Postepy Biologii Komorki, 47(3), 321–334.

Karamad, D., Khosravi-Darani, K., Hosseini, H., & Tavasoli, S. (2019). Analytical procedures and methods validation for oxalate content estimation. Biointerface Research in Applied Chemistry, 9(5), 4305.

Kavi Kishor, P. B., Anil Kumar, S., Naravula, J., Hima Kumari, P., Kummari, D., Guddimalli, R., Edupuganti, S., Karumanchi, A. R., Venkatachalam, P., & Suravajhala, P. (2021). Improvement of small seed for big nutritional feed. Physiology and Molecular Biology of Plants, 1–14.

Kavitha, S., & Sujatha, R. (2017). Iranian Journal of Neurology © 2017 Corresponding Author: http://ijnl.tums.ac.ir 4 April Atherogenic indices in stroke patients: A retrospective study. In Iran J Neurol (Vol. 16, Issue 2). http://ijnl.tums.ac.ir

Khan, S., Rubab, Z., Ali, I., Arshad, R., Abbas, A., & Akhtar, E. (2021). The Serum Electrolyte Imbalance in Children with Severe Acute Malnutrition. Journal of Islamic International Medical College (JIIMC), 16(1), 10–13.

Lambe, M. O. (2019). Ameliorative Effects of Moringa oleifera Leaf-based diet on Malnutrition-induced Skeletal Muscle Degeneration in Rats. DEPARTMENT OF BIOCHEMISTRY, FACULTY OF LIFE SCIENCES, UNIVERSITY OF ILORIN.

Levy, G. B. (1981). Determination of sodium with ion-selective electrodes. Clinical Chemistry, 27(8), 1435–1438.

Mathewson, S. L., Azevedo, P. S., Gordon, A. L., Phillips, B. E., & Greig, C. A. (2021). Overcoming protein-energy malnutrition in older adults in the residential care setting: A narrative review of causes and interventions. Ageing Research Reviews, 70, 101401.

Mendez, J., Franklin, B., & Gahagan, H. (1975). Simple manual procedure for determination of serum triglycerides. Clinical Chemistry, 21(6), 768–770.

Mertz, E. T., Hassen, M. M., Cairns-Whittern, C., Kirleis, A. W., Tu, L., & Axtell, J. D. (1984). Pepsin digestibility of proteins in sorghum and other major cereals. Proceedings of the National Academy of Sciences, 81(1), 1–2.

Mhlomi, Y. N., Unuofin, J. O., Otunola, G. A., & Afolayan, A. J. (2022). Assessment of rats fed protein-deficient diets supplemented with Moringa Oleifera Leaf Meal. Current Research in Nutrition and Food Science Journal, 10(1), 45–55.

Michael, H., Amimo, J. O., Rajashekara, G., Saif, L. J., & Vlasova, A. N. (2022). Mechanisms of Kwashiorkor-Associated Immune Suppression: Insights From Human, Mouse, and Pig Studies. Frontiers in Immunology, 13(May), 1–19. 10.3389/fimmu.2022.826268

Mihrete, Y. (2019). Review on anti nutritional factors and their effect on mineral absorption. Acta Scientific Nutritional Health, 3(2), 84–89.

Morales, F., Montserrat-de la Paz, S., Leon, M. J., & Rivero-Pino, F. (2023). Effects of malnutrition on the immune system and infection and the role of nutritional strategies regarding improvements in children’s health status: A literature review. Nutrients, 16(1), 1.

Naheed, N., Ahmed, H., Rana, D., & Aslam, K. (2021). Nadia Naheed et al Please cite this article in press Nadia Naheed et al, Study To Determine The Protein Energy Malnutrition. Total Proteins And Electrophoresis Of Serum Protein., Indo Am. J. P. Sci, 08(1), 1806–1811. http://www.iajps.com

Obadoni, B. O., & Ochuko, P. O. (2002). Phytochemical studies and comparative efficacy of the crude extracts of some haemostatic plants in Edo and Delta States of Nigeria. Global Journal of Pure and Applied Sciences, 8(2), 203–208.

Ofori, E. K. (2023). Lipids and Lipoprotein Metabolism, Dyslipidemias, and Management. In Current Trends in the Diagnosis and Management of Metabolic Disorders (pp. 150–170). CRC Press.

Ohanenye, I. C., Ekezie, F.-G. C., Sarteshnizi, R. A., Boachie, R. T., Emenike, C. U., Sun, X., Nwachukwu, I. D., & Udenigwe, C. C. (2022). Legume seed protein digestibility as influenced by traditional and emerging physical processing technologies. Foods, 11(15), 2299.

Okhiria, A. (2007). The Prevalence of Protein Energy Malnutrition in Nigeria and Dietary Management _a Revised Study. http://repository.elizadeuniversity.edu.ng/handle/20.500.12398/573

Oladele, G., Oladele, C., Ahmed, G., Gara, P., Dirisu, E., Amao, E., & Babalola, A. (2022). Pediatrics & Therapeutics Relationship between the Nutritional Status of Under-Five Children and Family Function as Seen in the Under-Five Clinic of Federal Medical Centre Bida, North Central Nigeria. Pediatr Ther, 12(2), 1000436. 10.35248/2161-0665.22.12.436.Citation

Olarewaju, O. O., Fajinmi, O. O., Naidoo, K. K., Arthur, G. D., & Coopoosamy, R. M. (2022). A review of the medicinal plants with immune-boosting potential. Journal of Medicinal Plants for Economic Development, 6(1), 15.

Organization, W. H. (2020). The double burden of malnutrition: priority actions on ending childhood obesity. World Health Organization. Regional Office for South-East Asia.

Orlien, V., Aalaei, K., Poojary, M. M., Nielsen, D. S., Ahrné, L., & Carrascal, J. R. (2023). Effect of processing on in vitro digestibility (IVPD) of food proteins. Critical Reviews in Food Science and Nutrition, 63(16), 2790–2839.

Otemuyiwa, I. O., & Adewusi, S. R. A. (2013). Fatty Acid, Carotenoid and Tocopherol Content of Some Fast Foods from a Nigerian Eatery. Journal of Food and Nutrition Research, 1(5), 82–86. 10.12691/jfnr-1-5-1

Otsuji, S., Mizuno, K., Ito, S., Kawahara, S., & Kai, M. (1988). A new enzymatic approach for estimating total and direct bilirubin. Clinical Biochemistry, 21(1), 33–38.

Poitevin, E. (2016). Official methods for the determination of minerals and trace elements in infant formula and milk products: A Review. Journal of AOAC International, 99(1), 42–52.

Raj Rai, S., Bhattacharyya, C., Sarkar, A., Chakraborty, S., Sircar, E., Dutta, S., & Sengupta, R. (2021). Glutathione: Role in oxidative/nitrosative stress, antioxidant defense, and treatments. ChemistrySelect, 6(18), 4566–4590.

Rajendram, R., Preedy, V. R., & Patel, V. B. (2015). Branched chain amino acids in clinical nutrition.

Raphael, J. E., Atanu, F. O., Mohammed, S. S., & Abdulrahman, S. I. (2020). Anti-anaemic effects of Moringa oleifera seeds (Lam) in protein energy malnourished rats. GSC Biological and Pharmaceutical Sciences, 13(1), 197–202.

Rasheed, A., Firdos, U., Seemab, F., Rani, M., Hanif, K., Hanif, S., Siddique, S., & Zafar, M. (2023). Prevalence and Determinants of Protein-Energy Malnutrition (Pem) among Children Under 5 years of age in a Rural Community of Lahore. Biological and clinical sciences research journal, 2023(1), 493.

Reitman, S., & Frankel, S. (1957). A colorimetric method for the determination of serum glutamic oxalacetic and glutamic pyruvic transaminases. American Journal of Clinical Pathology, 28(1), 56–63.

Remesar, X., & Alemany, M. (2020). Dietary energy partition: The central role of glucose. International Journal of Molecular Sciences, 21(20), 7729.

Rose, A. (2019). Characterization of lipids and the protein co-products from various food sources using a one-step organic solvent extraction process. West Virginia University.

Saa, R. W., Fombang, E. N., Ndjantou, E. B., & Njintang, N. Y. (2019). Treatments and uses of Moringa oleifera seeds in human nutrition: A review. In Food Science and Nutrition (Vol. 7, Issue 6, pp. 1911–1919). Wiley-Blackwell. 10.1002/fsn3.1057

Sardabi, F., Azizi, M. H., Gavlighi, H. A., & Rashidinejad, A. (2022). Potential benefits of Moringa peregrina defatted seed: Effect of processing on nutritional and anti-nutritional properties, antioxidant capacity, in vitro digestibility of protein and starch, and inhibition of α-glucosidase and α-amylase enzymes. Food Chemistry Advances, 1, 100034.

Schönfeldt, H. C., & Hall, N. G. (2012). Dietary protein quality and malnutrition in Africa. British Journal of Nutrition, 108(S2), S69–S76.

Segwatibe, M. K., Cosa, S., & Bassey, K. (2023). Antioxidant and antimicrobial evaluations of Moringa oleifera Lam leaves extract and isolated compounds. Molecules, 28(2), 899.

Shahidi, F., Varatharajan, V., Oh, W. Y., & Peng, H. (2019). Phenolic compounds in agri-food by-products, their bioavailability and health effects. Food Bioact, 5(1), 57–119.

Shankar, A. H. (2020). Mineral deficiencies. In Hunter’s Tropical Medicine and Emerging Infectious Diseases (pp. 1048–1054). Elsevier.

Shozib, H. B., Mahmud, S. A. S., Amin, R. B., Hossain, M. N., Biswas, J. K., Timsina, J., Ali, M. A., & Siddiquee, M. A. (2018). Nutraceutically enriched rice-based food to mitigate malnutrition in Bangladesh. EC Nutrition, 13(5), 240–249.

Shunmugapriya, K., Vennila, P., Thirukkumar, S., & Ilamaran, M. (2017). Identification of bioactive components in Moringa oleifera fruit by GC-MS. Journal of Pharmacognosy and Phytochemistry, 6(3), 748–751.

Siva Kiran, R. R., Madhu, G. M., & Satyanarayana, S. V. (2015). Spirulina in combating protein energy malnutrition (PEM) and protein energy wasting (PEW)-A review. J. Nutr. Res, 3(1), 62–79.

Soetan, K. O., & Aiyelaagbe, O. O. (2016). Proximate analysis, minerals, and anti-nutritional factors of Moringa oleifera leaves. Annals: Food Science & Technology, 17(1).

Sousa, R. M. L., Ribeiro, N. L. X., Pinto, B. A. S., Sanches, J. R., da Silva, M. U., Coêlho, C. F. F., França, L. M., de Figueiredo Neto, J. A., & Paes, A. M. de A. (2018). Long-term high-protein diet intake reverts weight gain and attenuates metabolic dysfunction on high-sucrose-fed adult rats. Nutrition & Metabolism, 15, 1–13.

Stechmiller, J. K. (2010). Understanding the role of nutrition and wound healing. Nutrition in Clinical Practice, 25(1), 61–68.

Steffes, M. W., & Freier, E. F. (1976). A simple and precise method of determining true sodium, potassium, and chloride concentrations in hyperlipemia. The Journal of Laboratory and Clinical Medicine, 88(4), 683–688.

Tahir, F., Ali, E., Hassan, S. A., Bhat, Z. F., Walayat, N., Nawaz, A., Khaneghah, A. M., Phimolsiripol, Y., Khan, M. R., & Aadil, R. M. (2024). Cyanogenic glucosides in plant-based foods: Occurrence, detection methods, and detoxification strategies–A comprehensive review. Microchemical Journal, 110065.

Tiloke, C., Anand, K., Gengan, R. M., & Chuturgoon, A. A. (2018). Moringa oleifera and their phytonanoparticles: Potential antiproliferative agents against cancer. Biomedicine & Pharmacotherapy, 108, 457–466.

Valarmathi, S., Pimpalshende, P. M., Seena, K. X., Arjun, U. V. N. V., Dasgupta, P., Mhaismale, V. K., & Mhaske, S. D. (2024). The Development And Assessment Of A Lotion Containing An Extract Of Moringa Oleifera L. Leaves With Varying Concentrations Of Cetyl Alcohol. Educational Administration: Theory and Practice, 30(4), 5569–5574.

Veniamin, M. P., & Vakirtzi-Lemonias, C. (1970). Chemical basis of the carbamidodiacetyl micromethod for estimation of urea, citrulline, and carbamyl derivatives. Clinical Chemistry, 16(1), 3–6.

Wei, Z., Zhou, N., Zou, L., Shi, Z., Dun, B., Ren, G., & Yao, Y. (2021). Soy protein alleviates malnutrition in weaning rats by regulating gut microbiota composition and serum metabolites. Frontiers in Nutrition, 8, 774203.

Wessels, I., Maywald, M., & Rink, L. (2017). Zinc as a gatekeeper of immune function. Nutrients, 9(12), 1286.

Wu, G. (2016). Dietary protein intake and human health. Food & Function, 7(3), 1251–1265.

Yakubu, M. T., Akanji, M. A., & Oladiji, A. T. (2005). Aphrodisiac potentials of the aqueous extract of Fadogia agrestis (Schweinf. Ex Hiern) stem in male albino rats. Asian Journal of Andrology, 7(4), 399–404. 10.1111/j.1745-7262.2005.00052.x

Zhang, J., Xu, X., Cao, Z., Wang, Y., Yang, H., Azarfar, A., & Li, S. (2019). Effect of different tannin sources on nutrient intake, digestibility, performance, nitrogen utilization, and blood parameters in dairy cows. Animals, 9(8), 507.

Zhang, Y., Hao, R., Chen, J., Li, S., Huang, K., Cao, H., Farag, M. A., Battino, M., Daglia, M., & Capanoglu, E. (2023). Health benefits of saponins and its mechanisms: perspectives from absorption, metabolism, and interaction with gut. Critical Reviews in Food Science and Nutrition, 1–22.

